# Two Chinese medicine species constants and the accurate identification of Chinese medicines

**DOI:** 10.1101/166140

**Authors:** Huabin zou

## Abstract

Since the ancient times, all over the world, the identification of herbal medicines have to be based on empirical knowledge. In this article two species constants of traditional Chinese medicines(TCM) were discovered relying on the maximum information states of Dual index information theory equation, or common heredity and variation information theory. The two species constants, common peak ratios *P*_g_ = 61% and *P*_g_ = 70%, which corresponding to symmetry and asymmetry variation states, respectively, were used as two absolute quantitative criteria to identify complex biology systems-TCM. Considered the influences of many other factors on components and experiment processes, the practical theoretical identification standards should be established *P*_g_≧58~64% and *P*_g_≧67~73%, within the relative error within −3% and + 3% of information value around the maximum information states. Combining the maximum number of effective sample optimum method with this two theoretical standards, the optimized classification of a TCM sample set can be carried out correctly. 42 samples belonging to four species of combination Chinese medicines were tested. The infrared (IR) fingerprint spectra (FPS) of their powder were measured and analyzed by means of the approach provided above. Among the six pairs of four Chinese medicine species, five of them follow the species constant *P*_g_=61%, one of them obeys the *P*_g_ = 70%. The correct recognition ratio of samples was 95.2%, and that of species was 100%.

## Introduction

The species of herbal medicines determine their efficacy, thus the accurate identification of species is very significant for herbal medicines, and for traditional Chinese medicines whose major part is composed of herbs and animals. In fact, the identification of Chinese medicines belongs to that of biology species. However, until recently there is no one unquestionably scientific definition or quantitatively standard relative to biology species[1,2,3]. This also leads to classification of complex organisms is in chaos which hampers biology conservation heavily presently [4]. Most importantly, in biology, species is the fundamental of biological diversity, and is the ultimate goal of systems biology [5]. In practical the classification or pattern recognition of Chinese medicines as well as biology is performed depending on some macro properties, such as macro characters, biological functions, geographic distributions, ecological environments and so forth. In the procedure, the criteria were established only rest on subjective deduce by researchers. Recently, all definitions about biology species lack of principle theory, were not based on experimental science and mathematical principles, and also are short of the quantitative standards. That is, the identification of biology and Chinese medicines are totally depending on their shapes, sizes, colors, tastes and the like, and empirical knowledge[6]. Even if in modern numerical taxonomy, the criteria are determined on the basis of a lot of learning-based experience.

As a kind of complex biology systems, whether TCMs are of some intrinsically accurate laws and pose of some precisely quantitative criteria, that is, some constants, like in physics and chemistry systems. Thus by applying some accurately quantitative criteria we can determine the TCM species precisely without experience or prior knowledge. This is a greatly significant scientific subject need to be investigated seriously. Author Zou has paid a great effort to searching for the mathematical theory for identifying traditional Chinese medicines and biological systems for long time, grounded on intrinsic properties and elemental principle in nature. He proposed and established independently the common and variant peak ratio dual index sequence analysis method during his doctorate work[7-13]. This theory has been adopted by many researchers to identify herbs, plants, foods and animals, etc, and more than 50 papers using this method have been published so far, such as papers [14-26]. Based on all these works, author Zou and coworkers established the dual index grade sequence individualized pattern recognition approach [27-34]. The Chines medicines’ species can be identified accurately relying on this method by selecting suitable similarity scales *P*_g_≧*P_g_* + *x S*, where −3≦*x*≦3, without any help of empirical knowledge and learning samples. In the procedure, by change *x* smoothly to obtain each sample’s characteristic sequence. Samples in core characteristic sequences of each class construct an independent set, and different sets represent different classes. However, this approach can’t offer any absolute quantitative criterion for Chines medicine species.

In paper [35] a theory—the Dual index information theory or heredity and variation information theory equation was proposed. According to this theory, there are two intrinsically variation mechanism, which are symmetry and asymmetry variations. Based on the two variation mechanisms, two common peak ratios *P*_g_=0.61 and *P*_g_=0.70, corresponding to the maximum information states, are defined as two constants, in which the constant related to symmetry variation was proved to fit to classify some combination Chinese medicines [35,36,37] successfully, by means of standard *P*_g_≧61%. These complex biology systems include components extracted from Guifu Dihuang Pills, Jinkui Shenqi Pills with absolute ethanol[35], the powder of Mingmu Dihuang Pills, Zhibai Dhuang Pills, and Maiwei Dihuang Pills[36], the components of Buzhong Yiqi Pills and Shiquan Dabu Pills extracted with chloroform[37]. While there is no report about the other standard *P*_g_≧70% applied to identification biology systems. This article focuses on to investigate the relationship between Chinese medicine species pairs, in order to find whether some Chinese medicines species obeythe two absolute quantitative criteria, especially the constant *P*_g_≧70%. In this paper, 6 pairwise of four species of Chinese medicines: Guifu Dihuang Pills, Jinkui Shenqi Pills, Mingmu Dihuang Pills, Zhibai Dihuang Pills were investigated subtly. The infrared fingerprint spectra of 42 samples’ powders belonging to the four species were measured and analyzed. Compared to components extracted from Chinese medicines with some kind of solvent, compounds existed in Chinese medicine powder can fully reflect their properties or features. The Infrared fingerprint spectra of their powder are better than that of their extracted components for verifying theory. All six pairwise of these four Chinese medicines were tested, the results showed among them, five pairs were successfully classified based on *P*_g_≧61%, and only one pair Guifu Dihuang Pills and Zhibai Dihuang Pills obeys the criterion *P*_g_≧70%. These 42 samples and the six Chinese medicine pairs were distinguished perfectly. The accuracy of sample recognition was 95.2%. These results proved that there really exist the two constants in some complex organisms, and they can be defined as Chinese medicine species constants, which exhibit the intrinsic quality characteristic. Depending on these two Chinese medicine species constants, some biology systems can be precisely identified, pattern recognized, without help of any prior knowledge relative to sample set.

The Infrared fingerprint spectra of these 42 samples’ powder were also analyzed by means of the dual index grade sequence individualized pattern recognition method [27-34]. When selected the similarity scale to be *P*_g_≧*P_g_* + 1.1*S*and *P*_g_≧*P_g_* + 1.3*S*, these 42 samples were accurately classified with the correct ratio being 90.5% and 95.2%, respectively. The results gained rest on the two theoretical approaches are equivalent to each other. While the two Chinese medicine species constants are the absolute quantitative criteria, they make the identification of biology systems and Chinese medicines being of the most simplicity, clear physics and biology meanings. On the other hand, this work, to put forward two Chinese medicine species constants may be a breakthrough from ancient times.

## 1 Two TCM species constants

According to the Dual index information theory equation-the heredity and variation information theory equation[35], the information of any two biological samples or any two evolution stages of one biological system can be calculated by means of the follow equation:

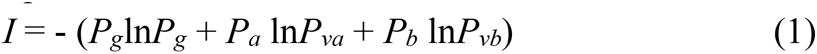

The meaning of every parameters /variables and its definition are showed as follows, as in paper[6-10].

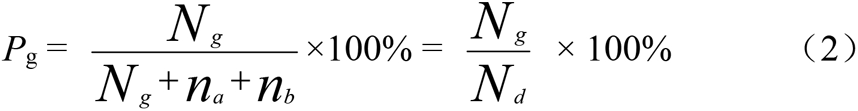

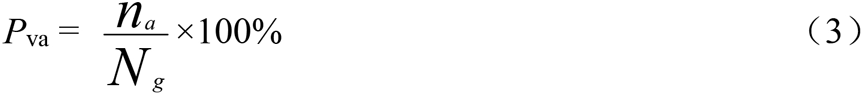

Here *P*_g_ is common peak ratio of fingerprint spectra *a* and *b*. Its value is the ratio of number of common peaks *N _g_* to that of total kinds of peaks *N_d_* in *a* and *b*, *N_d_* is defined as independent peaks, that is, *N_d_* is the kinds of peaks. *N _g_* are the peaks existed both in fingerprint spectra *a* and *b*.

Interestingly, this parameter is the same as Jaccard parameter, Sneath and Sokal parameter[38]. *n_a_* and *n_b_* are the number of variation peaks only existed in fingerprint spectra *a* and *b*, respectively.

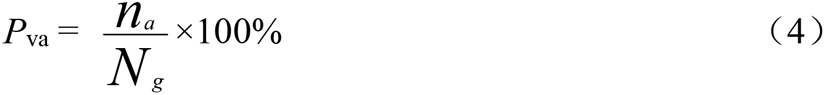

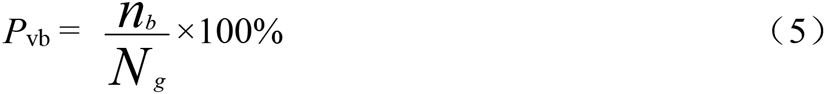

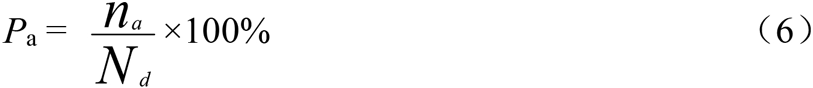

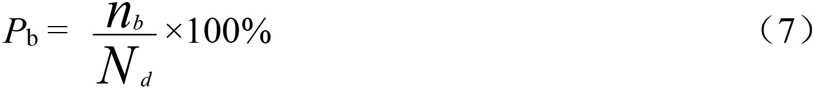

*P_a_* is the existed probability or ratio of *n_a_* to *N_d_*. *P_b_* is that of *n_b_* to *N_d_ P _va_* and *P_vb_* are defined as variation peak ratio of *n_a_* and *n_b_* to *N _g_*, respectively.

The relationships among different peaks are listed here.

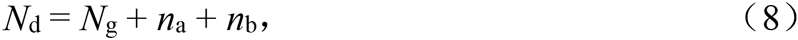

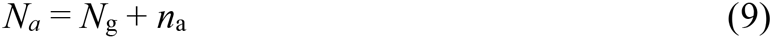

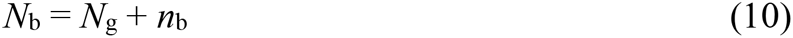

In order to describe the symmetric degree of among *n_a_* and *n_b_*, that is among system *a* and *b*, a new parameter—the symmetric degree αis defined as 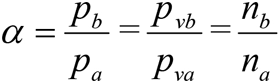, 0≦α≦1. When α=1, It means perfectly symmetric variation, and α=0, It means extremely asymmetric variation. There are two maximum information values corresponding to the two states, the α=1, *P_g_* =0.61, and α=0, *P_g_* =0.70. These states can be displayed in figure 1.

**Fig. 1.**
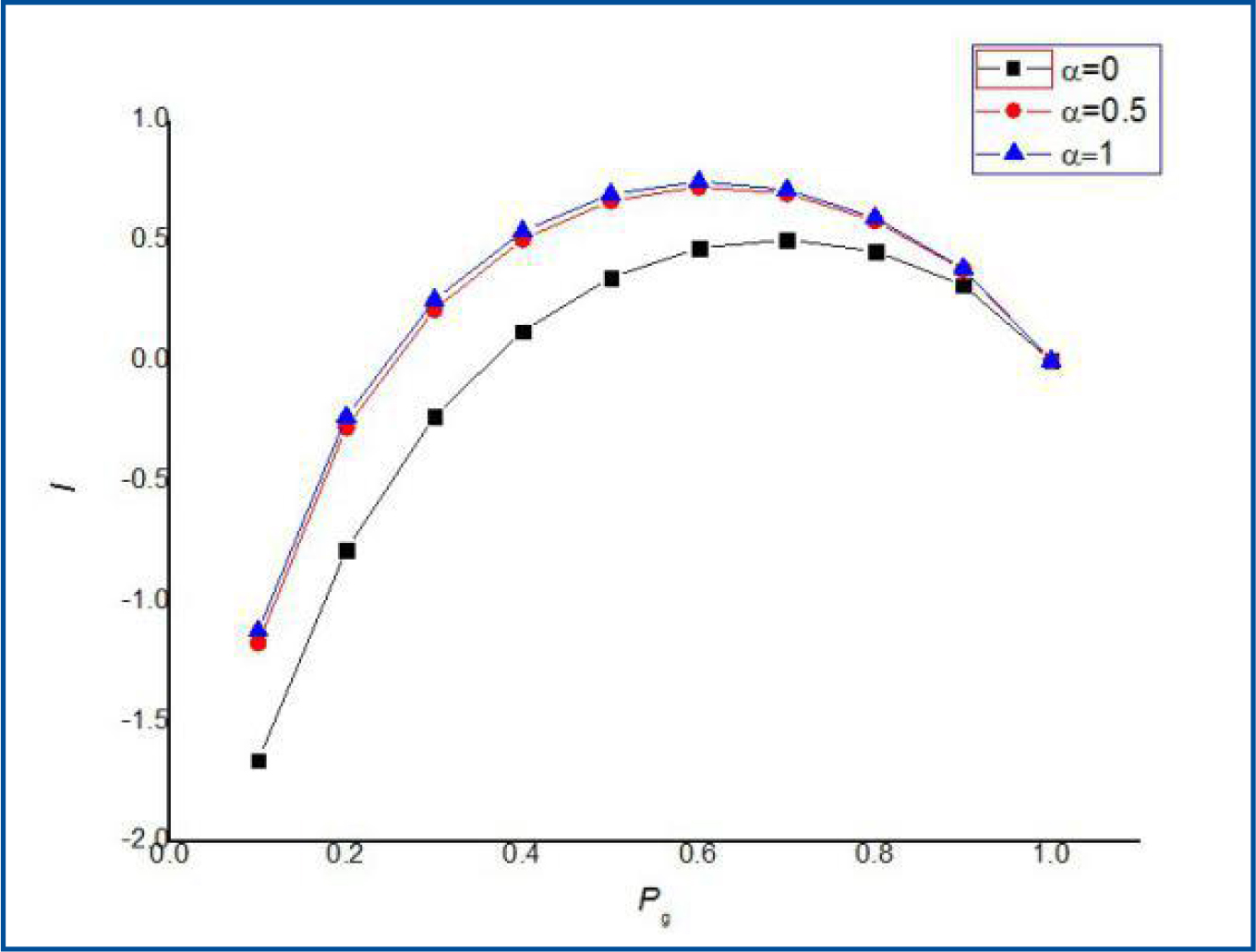
the *I* ~ *P_g_*curves of dual index information equation.

In accordance with *I*~*P*_g_ figure, one can find that the top points of *I* are at *P*_g_=0.61=61%, when α= 0.5, 1. While that of *I* is at *P*_g_=0.695 = 0.70 =70%, when α = 0. Moreover, there is an flat interval existed near *P*_g_=0.61 and 0.70. In each interval *I* changes little. *P*_g_ corresponding to the maximum information intervals, and the change ratios of information corresponding to these *P*_g_ intervals, at different α, are shown in table 1.

**Table 1.**
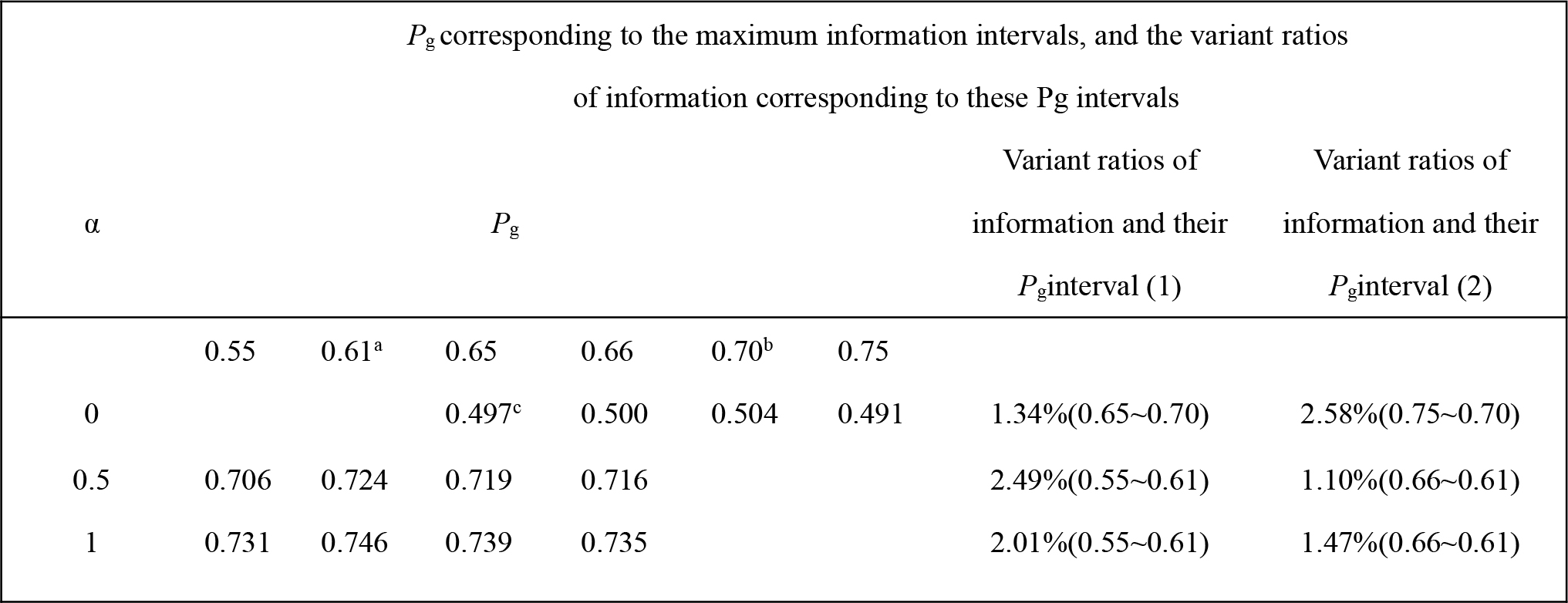
*P*_g_ corresponding to the maximum information intervals, and the change ratios of information corresponding to these *P*_g_ intervals at different α

| $P_g$ corresponding to the maximum information intervals, and the variant ratios of information corresponding to these $P_g$ intervals |  |  |  |  |  |  |  |  |
| --- | --- | --- | --- | --- | --- | --- | --- | --- |
| $\alpha$ | $P_g$ | | | | | | Variant ratios of | Variant ratios of |
|  |  |  |  |  |  |  | information and their | information and their |
| | | | | | | | $P_g$interval (1) | $P_g$interval (2) |
|  | 0.55 | 0.61<sup>a</sup> | 0.65 | 0.66 | 0.70<sup>b</sup> | 0.75 |  |  |
| 0 |  |  | 0.497<sup>c</sup> | 0.500 | 0.504 | 0.491 | 1.34%(0.65~0.70) | 2.58%(0.75~0.70) |
| 0.5 | 0.706 | 0.724 | 0.719 | 0.716 |  |  | 2.49%(0.55~0.61) | 1.10%(0.66~0.61) |
| 1 | 0.731 | 0.746 | 0.739 | 0.735 |  |  | 2.01%(0.55~0.61) | 1.47%(0.66~0.61) |

*a*, when symmetric degree α= 0.5, 1, the maximum information corresponding to *P*_g_ = 0.61; *b*, when α=0, the maximum information is at *P*_g_=0.70. *c*, the information values corresponding to *P*_g_ upper.

The theoretical analysis offered above point out, when α= 0.5~1, the maximum information are all at *P*_g_=0.61. When the *P*_g_ interval region from 0.55 to 0.66, the information change little, only within ±3%. When α=0, the maximum information is at *P*_g_=0.70, and when *P*_g_ interval region is from 0.65 to 0.75, the information also varies within ±3%. These results indicated when *P*_g_ changes within ±5%, from *P*_g_ corresponding to the maximum information, the *I* vary within ± 3%. This region may be considered as the critical characteristic zone of TCM species.

The higher the common peak ratio between samples, the larger the similarity between them. This means their quality is more similar to each other. Therefore, relying on table 1, *P*_g_=61% and *P*_g_=70% can be regarded as or defined as two TCM species constants. Depending on the two absolute theory criteria, wen can reasonably set practical theory standards *P*_g_≧(61 ± 3)% and *P*_g_≧(70 ± 3)% as two optimizing and judging TCM species, considering all kinds of factors affecting experiment results.

## 2 material

### 2.1 Regents and Chinese medicine samples

KBr (AR) were purchased from Tianjing national regent company (China), and 42 samples belonging to four species of combination Chinese medicines: Guifu Dihuang pills, Jinkui Shenqi Pills, Mingmu Dihuang Pills and Zhibai Dihuang Pills are shown in Table 2.

**Table 2.**
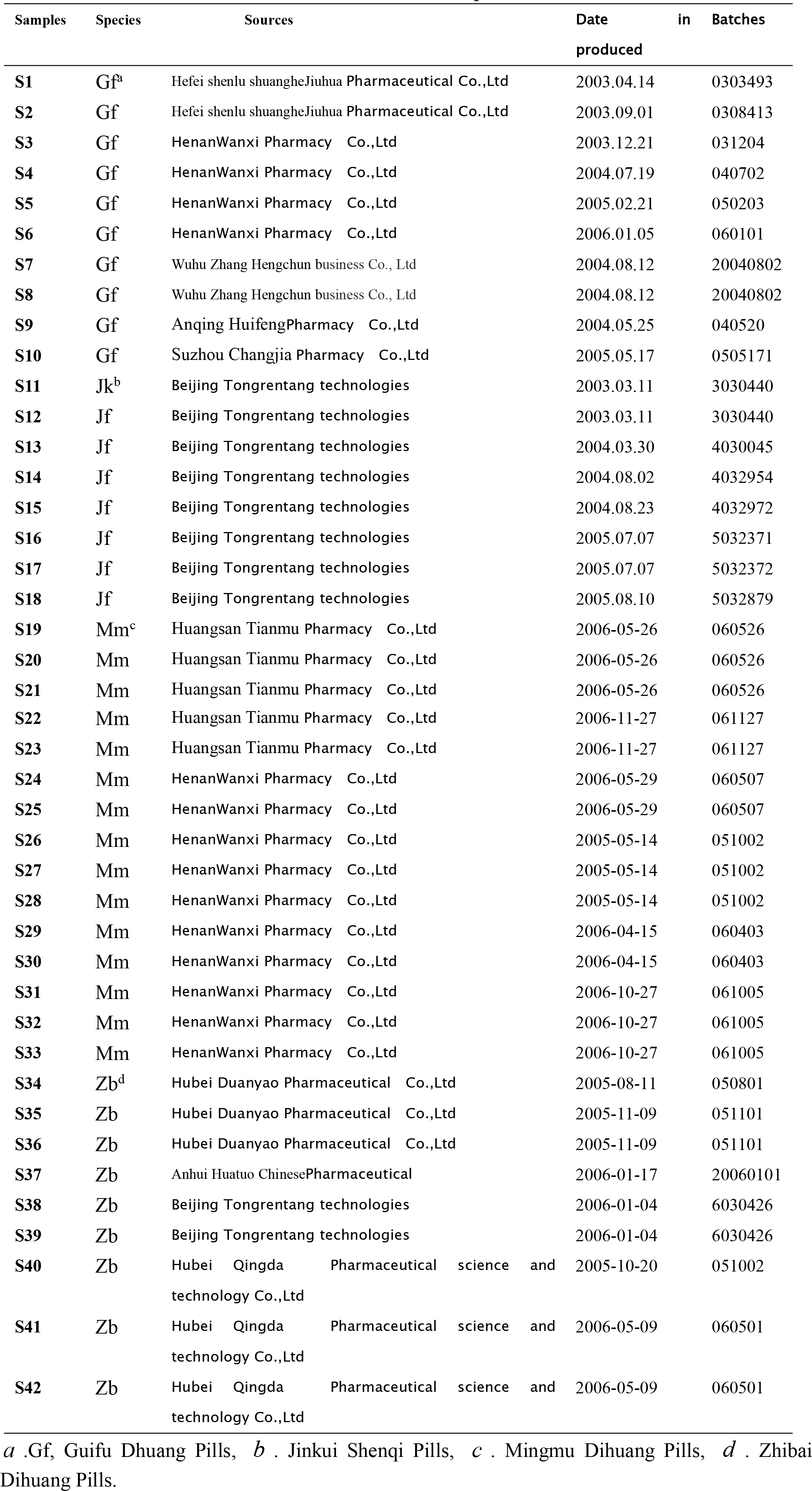
*P* The sources of samples of TCM

### 2.2 Composition of four TCM species [39]

The composition of four species of TCM investigated in this paper are as follows.

**Guifu Dhuang Pills:** Radix Rehmanniae, Poria, Cortex Moutan, Rhizoma Dioscoreae, Fructus Corni, Rhizoma Alismatis, Radix Aconiti Later alis Preparata, Cortex Cinnamomi.

**Jinkui Shenqi Pills:** Radix Rehmanniae, Poria, Cortex Moutan, Rhizoma Dioscoreae, Fructus Corni, Rhizoma Alismatis, Radix Aconiti Later alis Preparata, Cortex Cinnamomi, Radix Achyranthis Bidentatae, Plantain seed, Cassia twig.

**Mingmu Dihuang Pills:** Radix Rehmanniae, Poria, Cortex Moutan, Rhizoma Dioscoreae, Fructus Corni, Rhizoma Alismatis, Radix Paeoniae Alba, Radix Angelicae Sinensis, Fructus Lycii, Fructus Tribuli, Flos Chrysanthemi, Concha Haliotidis.

**Zhibai Dihuang Pills:** Radix Rehmanniae, Poria, Cortex Moutan, Rhizoma Dioscoreae, Fructus Corni, Rhizoma Alismatis, Cortex Phellodendri, Rhizoma Anemarrhenae.

**The list** showed all four species of TCM contain common six kinds of herbs, Radix Rehmanniae, Poria, Cortex Moutan, Rhizoma Dioscoreae, Fructus Corni, Rhizoma

Alismatis. Thus these four species of TCM are composed of a great deal of the same or similar compounds, which make it very difficult to accurately identify them or classify them.

## 3 Results and discussion

### 3.1 Detection of infrared fingerprint spectra

In keeping with the experiment conditions listed in methods, the infrared fingerprint spectra of 42 samples’ powder belonging to four species of traditional Chinese medicines, were measured, showed in figure 1~4. From these figures, one can notice they are very analogous, and are of high complexity. It is difficult to distinguish them by direct observation. Their accurate identification must be carried out depending on mathematical theory to analyze these infrared fingerprint spectra.

**Fig. 1.**
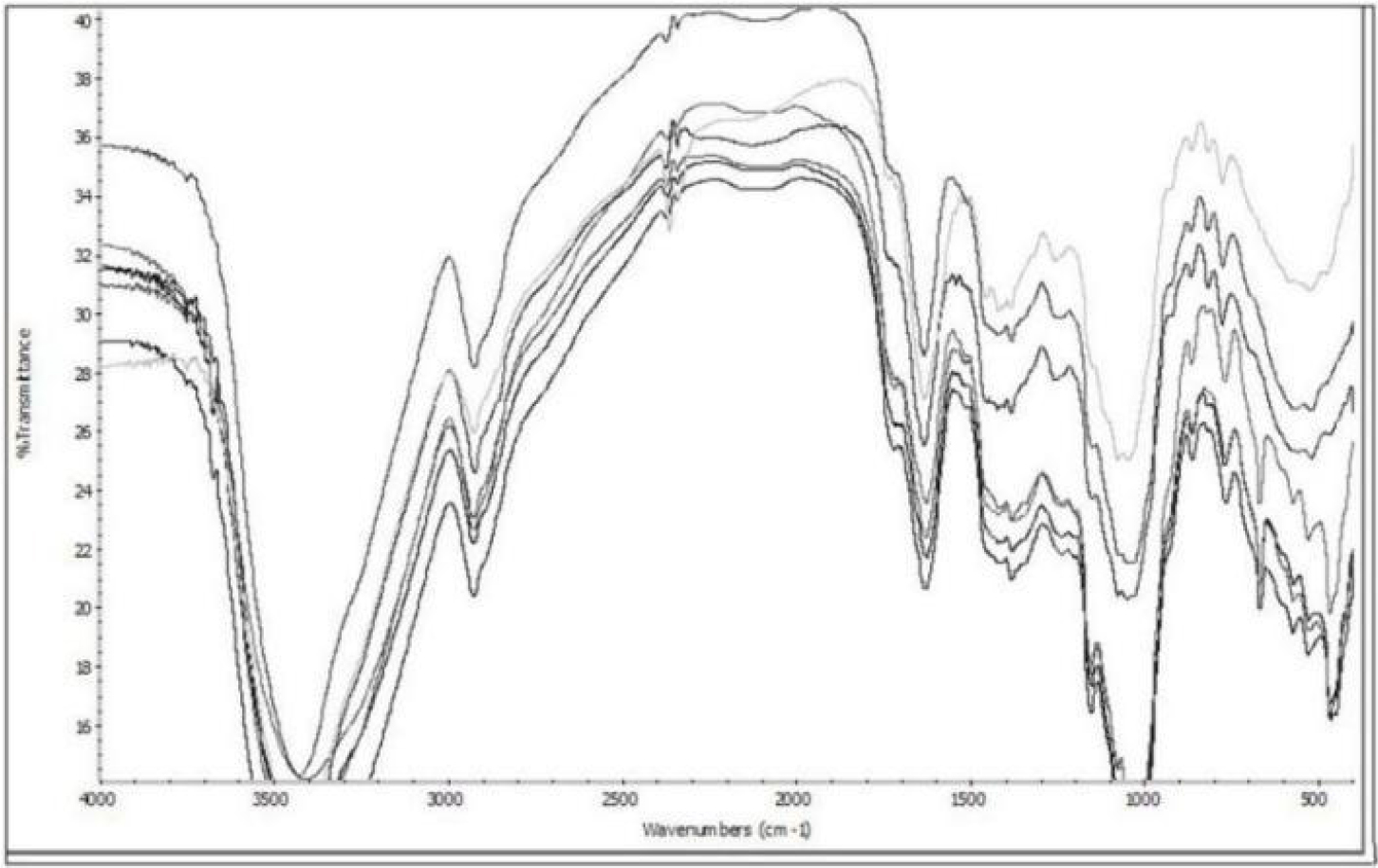
the overlapped infrared fingerprint spectra of Guifu Dihuang pills and Jingkui Shenqi Pills’ powders. They were S1,S3,S8,S10 (Guifu Dihuang pills), S11,S14,S18 (Jingkui Shenqi Pills) at near 1600cm^−1^ from bottom to top.

**Fig. 2.**
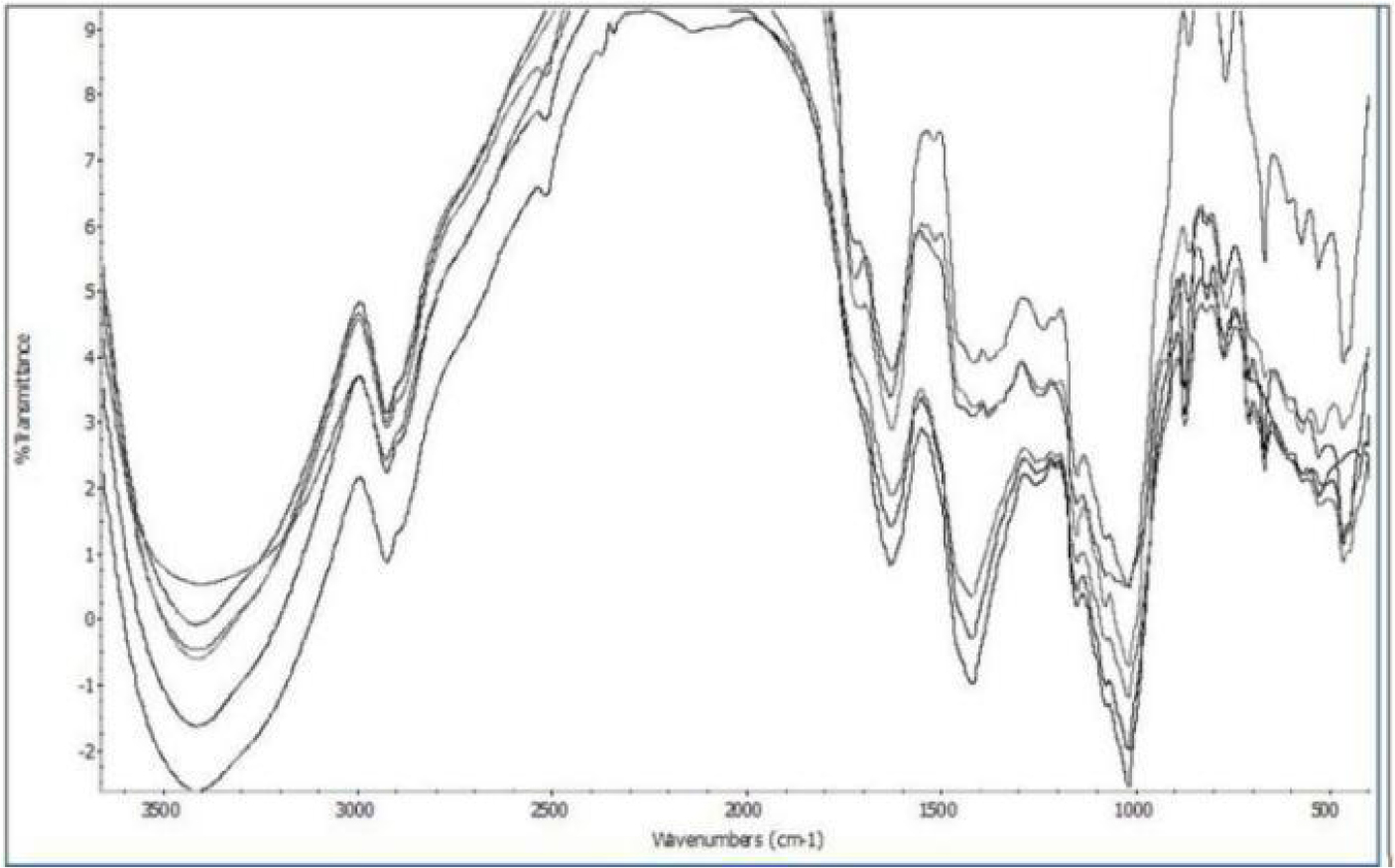
the overlapped infrared fingerprint spectra of Mingmu Dihuang pills and Zhibai Dihuang Pills’ powders. They were S24,S26,S32(Mingmu Dihuang pills), S34,S38,S42 (Zhibai Dihuang Pills) at near 1600cm^−1^ from bottom to top.

**Fig. 3.**
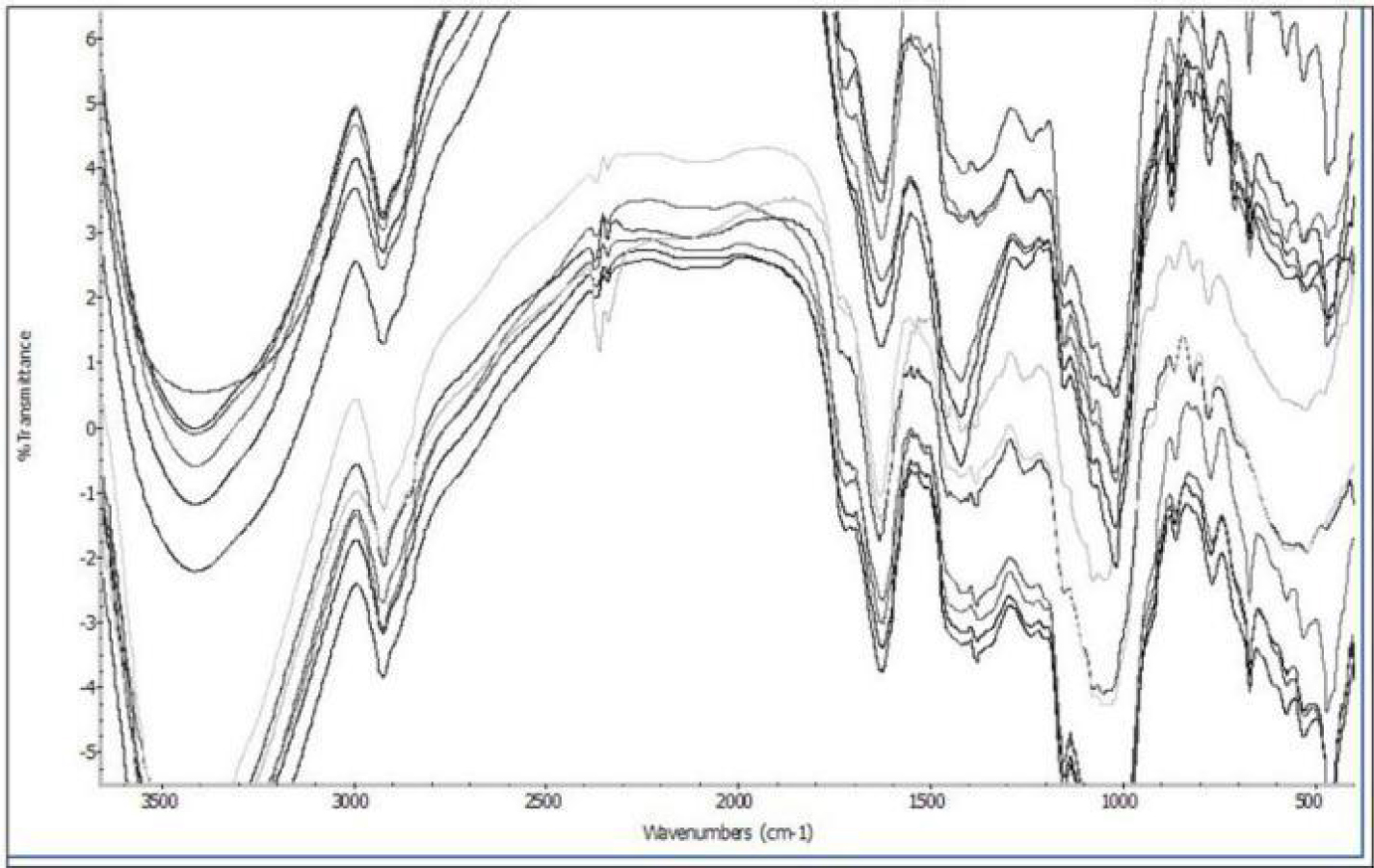
the overlapped infrared fingerprint spectra of Guifu Dihuang pills, Jingkui Shenqi Pills, Mingmu Dihuang pills and Zhibai Dihuang Pills’ powders. They were S1,S3,S8,S10(Guifu Dihuang pills), S11,S14,S18 (Jingkui Shenqi Pills), S24,S26,S32 (Mingmu Dihuang pills), S34,S38,S42(Zhibai Dihuang Pills), at near 1600cm^−1^ from bottom to top.

**Fig. 4.**
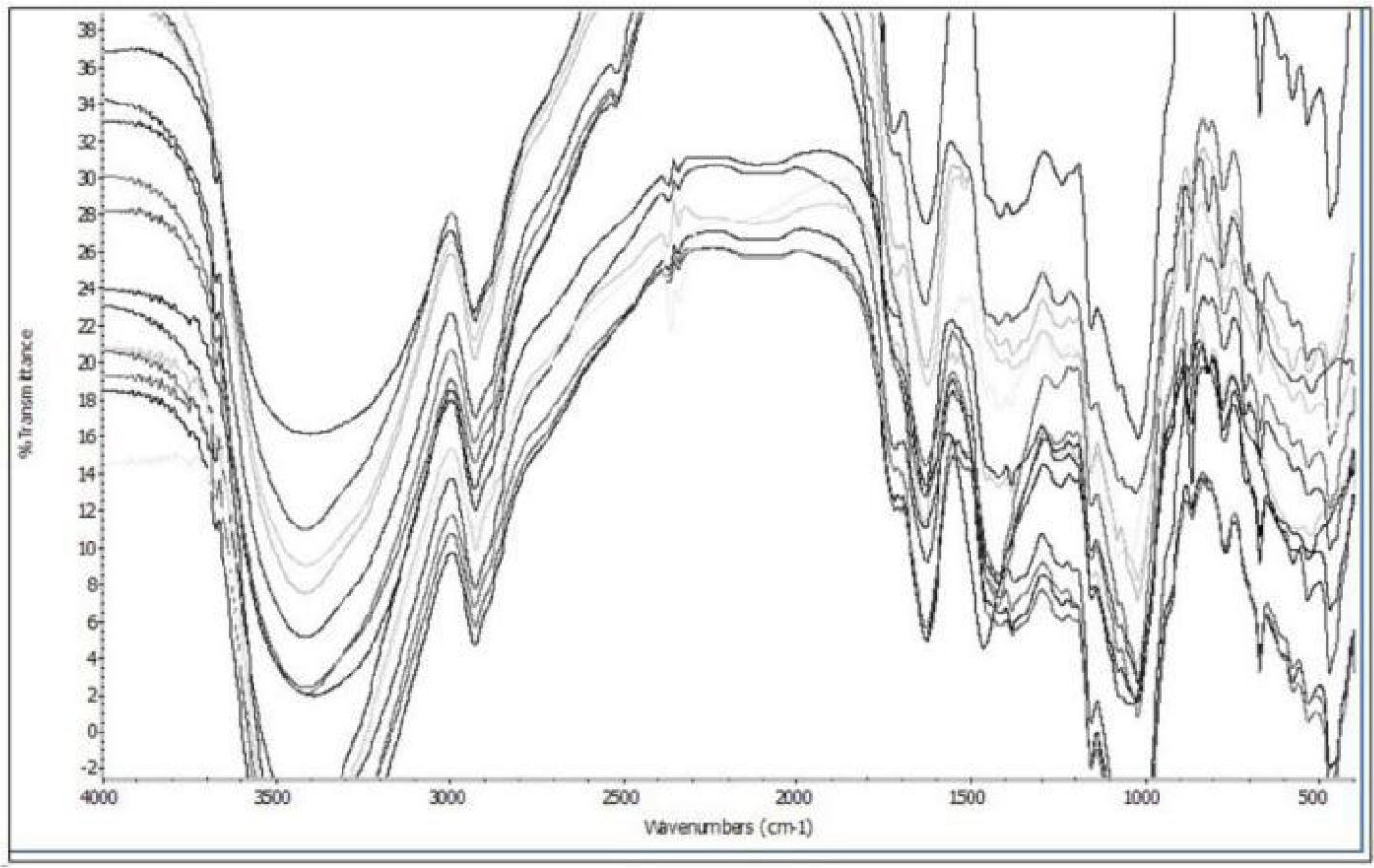
the overlapped infrared fingerprint spectra of Guifu Dihuang pills, Jingkui Shenqi Pills, Mingmu Dihuang pills and Zhibai Dihuang Pills’ powders. They were S1,S3,S8,S10 (Guifu Dihuang pills), S11,S14,S18 (Jingkui Shenqi Pills), S19,S24,S26,S32 (Mingmu Dihuang pills), S34,S37,S38,S42 (Zhibai Dihuang Pills), at near 2900cm^−1^ from bottom to top.

### 3.2 Analysis on data

To determine common and variant peaks of infrared fingerprint spectra of these 42 samples based on the Shapiro-Wilk W-testing method[40]. To select any sample as a reference, and calculate the common peak ratios of the rest samples to the reference. Then to place these rest sample marks, together with their common peak ratios in the order: common peak ratio from high to low to form a sequence.

Based on the two traditional Chinese medicine species constants *P*_g_ = 0.61 and *P*_g_ = 0.70, and combining the maximum number of effective samples method in characteristic sequence of sample set [36].

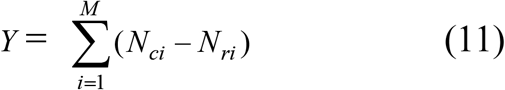

*Y*: the maximum effective samples in characteristic sequences of sample set.

*N_ci_*: the number of samples in the *i* th samples’ characteristic sequence.

*N_ri_*: the number of samples in *i* th samples’ related sequence.

*M*: the number of total samples.

*Y* reflects degree of effective classification of sample set. The larger the *Y*, the more clear the classification, and the much less the samples in related sequences. This means that the more samples in core characteristic sequences, the classification is more ideal.

To optimizeclassification relying on *P*_g_≧(61±3)% and *P*_g_≧(70±3)%, and to get integrally optimized characteristic sequences. Then to carried out the concisely identification and classification of these 42 samples.

When *P*_g_≧61% and *P*_g_≧72%, the optimized characteristic sequences of the 42 samples belonging to four TCM species were showed as follows.

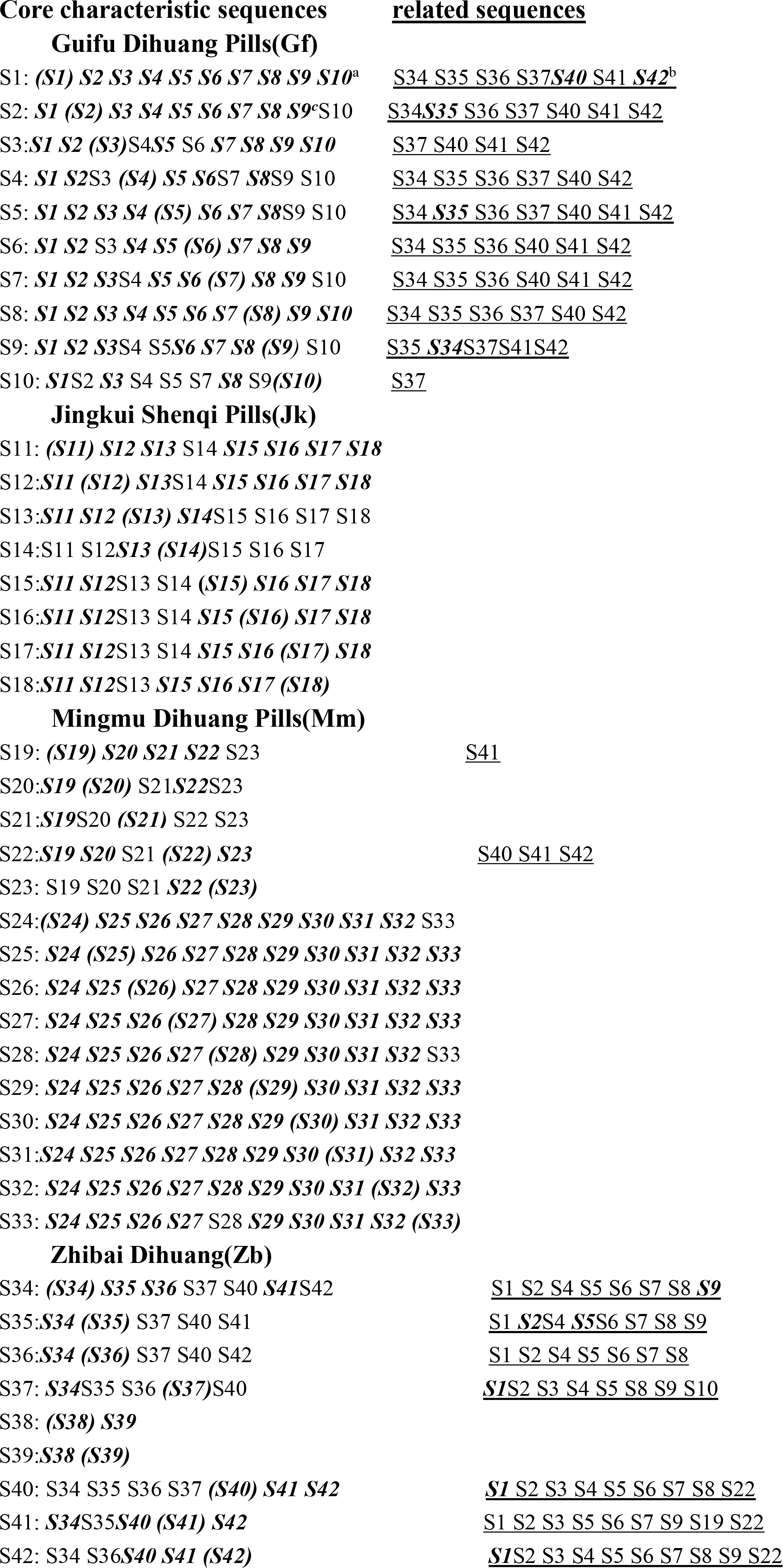

*a*, the core characteristic sequences; *b*, the related sequences, samples in this part not belonging to the class to that the samples in core characteristic sequences corresponded. characteristic sequences is made up of core characteristic sequences and related sequences. *c*, the samples represented by ***black-itaics*** are the characteristic sequences of 42 samples, when choose *P*_g_≧72%. While the whole sequences are the characteristic sequences when *P*_g_≧61%.

When *P*_g_≧61%, five of the six pairwise, constructed from four TCM species, could be classified accurately according to each samples’ characteristic, listed ahead. They were Gf-Jk, Gf-Mm, Jk-Mm, Jk-Zb, Mm-Zb. While there existed heavy overlap betwee Gf-Zb smaples’ characteristic sequences exhibited above. This indicated when *P*_g_≧61%, Gf and Zb samples could not be classified.

When *P*_g_≧72%, for S35,S37, samples in related sequences are more than that in core characteristic sequences. Thus S35, S37 could not be corrected recognition. However, other 40 samples could be identified clearly. The correct ratio was 40/42= 95.2%. Under this condition, all six pairs were identified and classified perfectly. Based on the TCM species constant *P*_g_=70%, together with the maximum effective sample optimum method[34], to change *P*_g_ according to *P*_g_≧(70± *x*) %, *x* =-1, 0, 1, 2, 5, the number of maximum effective samples were listed in table 3.

**Table 3.** The number of effective samples based on *P*_g_≧(70± *x*) %

| $P_g\%$ | Optimized results | |
| --- | --- | --- |
|  | The maximum number<br>of effective samples | Correct recognition<br>ratio (%) |
| $\geq 69$ | 249 | 92.9 |
| $\geq 70$ | 254 | 92.9 |
| $\geq 71$ | 254 | 92.9 |
| $\geq 72$ | **256** | 95.2 |
| $\geq 75$ | 243 | 100 |

$x^*$ , $x = -1, 0, 1, 2, 5$ .

From table 3, when *P*_g_≧69~75%, all samples could be distinguished completely. However, when *P*_g_≧72%, the number of the effective samples was the largest one. The result was most reasonable. Further, when *P*_g_≧75%, the correct ratio was 100%, which is larger than 95.2%, but the number of the effective samples was the lowest, only was 243. In this case, the over classification was carried out, the results were not reasonable.

Analysis on the data from the IR FPS of 42 sample powders, belonging to four TCM species, showed that the identification criteria of these four TCM species are *P*_g_≧61% and *P*_g_≧75%, the results were listed in table 4.

**Table 4.**
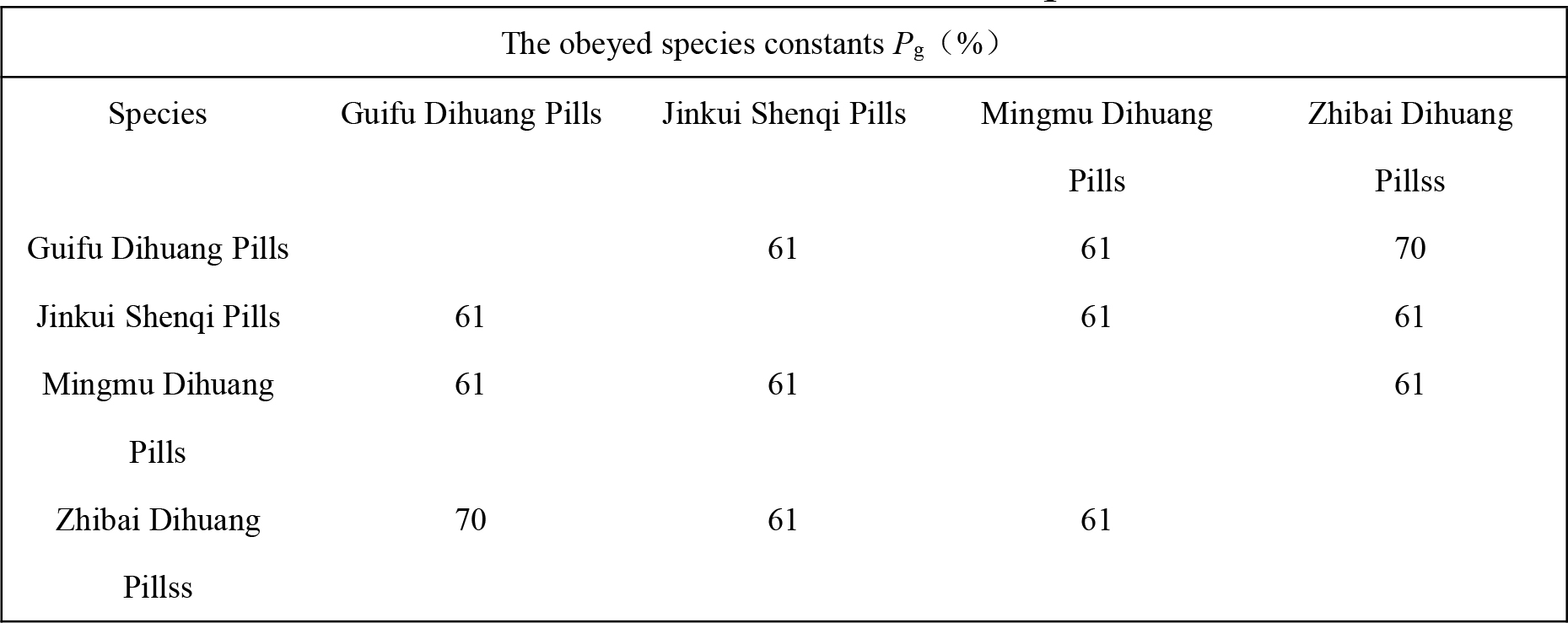
test results of the two TCM species constants

Under the similarity scales showed in table 4, all six pairs among four TCM species were classified correctly. The correct ratio of samples was 95.2%. In this case, the number of effective samples was the largest. The correct ratio of TCM species was 100%.

On the other hand, the IR FPS could be analyzed by means of the dual index grade sequence individualized pattern recognition method. When similarity scale is *P_g_*≧*P_g_* + 1.3*S*, S35 was not classified successfully. S37 could not be recognized because the number of samples in its related sequence is equal to that in its core characteristic sequence. The correct recognition ratio is 40/42 = 95.2%, which is the same as that obtained by means of the Two TCM species constants. These results proved that this new method is reliable.

## 4 conclusion

Depending on the two TCM species constants *P*_g_=61% and *P*_g_=70%, obtained from the dual index information theory equation, and on the two theoretical standards *P*_g_≧(61 ± 3)% and *P*_g_≧(70 ± 3)%, established rest on the two TCM species constants, 42 samples and the six pairs combined from four TCM species were perfectly identified and classified, together with the maximum number of effective samples method [36]. These results can support conclusion that the two common peak ratios deduced from the dual index information theory equation, corresponding two maximum information states, which related to two extremely variation states, are able to truly reflect the intrinsic characteristic of complex biology systems. This theory could concisely identify, classify some complex biology systems without any help of prior knowledge, such as some learning samples, some parameters decided subjectively about sample set. These work were achieved only relying on the measured data from sample set and the maximum number of effective sample method. In this approach, the maximum number of effective samples enable researchers to avoid over classifying a sample set, and to achieve an optimum pattern recognition result. This verifies that in biology systems, there exist some precisely science laws, and some constants, like in physics and chemistry. These original work will attract other researchers to discover some precisely science laws in biological systems in the future.

Trace back to several thousand years, it may be a great breakthrough by far to accurately identify TCM species, or biology systems by means of absolute quantitatively theoretical criteria, not experience, prior knowledge. Moreover, this theory is of great simplicity, and may be suit to other biology systems in principle.

## 5 Methods

### 5.1 Instruments

FT-IR spectrophotometer ModelNICOLET-5700-FT-IR(USA), withspectral range: 4000–400 cm^−1^, resolving power 4 cm^−1^; high speed grinder; tablet press; analytical balance were used in this study.

### 5.2 Conditions

All samples were powdered. To shift the powders with 80 eye sieve, then to heat them at 60°C for 2 hours, and keep them at below 4°C. The experiments show if the powders are kept at low temperature, the IR FPSobtained remain the same and no degradation is observed.

The infrared fingerprint spectra of these samples were measured by means of KBr tablettingmethod. Each sample was detected for 6 times to obtain 6 infrared fingerprint spectra, based on which a combination numerical fingerprint spectrum was formed. The wavenumbers of each peak in the combination numerical fingerprint spectrum were the mean of that of this peak appeared in the 6 infrared fingerprint spectra. These infrared fingerprint spectrawere smoothed by 25 cm^−1^at sensitivity 80.

### 5.3 Repeatability and stability

The sample S20 was parallelly measured for 6 times and got 6 infrared fingerprint spectra, and choose any four fingerprint spectra to construct a combination numerical fingerprint spectra, in which every peak’ s wavenumbers is the average of the peaks appeared in the same position in four fingerprint spectra. The results showed the minmum *P*_g_ was 93% of among these combination numerical fingerprint spectra. Thus the fingerprint spectra were of excellent repeatability. The powders of traditional Chinese medicines were kept below 4°C, and were of good stability.

